# Molecular Dynamics Analysis of Mg^2+^-dependent Cleavage of a Pistol Ribozyme Reveals a Fail-safe Secondary Ion for Catalysis

**DOI:** 10.1101/596270

**Authors:** Newlyn N. Joseph, Raktim N. Roy, Thomas A. Steitz

## Abstract

Pistol ribozymes comprise a class of small, self-cleaving RNAs discovered via comparative genomic analysis. Prior work in the field has probed the kinetics of the cleavage reaction, as well as the influence of various metal ion cofactors that accelerate the process. In the current study we perform unbiased and unconstrained molecular dynamics simulations from two current high-resolution pistol crystal structures, and we analyzed trajectory data within the context of the currently accepted ribozyme mechanistic framework. Root-mean-squared deviations (RMSDs), radial distribution functions (RDFs), and distributions of nucleophilic angle-of-attack reveal insights into the potential roles of three magnesium ions with respect to catalysis and overall conformational stability of the molecule. A series of simulation trajectories containing *in-silico* mutations reveal the relatively flexible and partially interchangeable roles of two particular magnesium ions within solvated hydrogen-bonding distances from the catalytic center.

## I. INTRODUCTION

Currently fourteen well-defined naturally-occurring ribozyme classes are known to exist [1–8]. Nine among these are self-cleaving and have been linked to several distinct biological functions [2, 5, 6]. These small structured RNA molecules employ various mechanistic strategies to achieve scission of the phosphodiester linkage between two adjacent nucleotides [9, 10]. The reaction is characterized by a nucleophilic attack of a 2′-oxygen atom on the adjacent phosphorus atom [7, 8, 10–13]. This process liberates the 5′ segment and results in the formation of a 2′,3′-cyclic phosphate. This cleavage reaction undergoes a trigonal-bipyramidal transition state, which can benefit from stabilization of the strained geometry and neutralization of the developing charges to obtain better reaction rates [10, 14, 15]. Ribozymes are believed to promote catalysis by utilizing multiple catalytic strategies drawn from the following list: in-line attack, stabilization of the negative charge on the non-bridging phosphate oxygen atoms, proton abstraction from the 2′-hydroxyl and stabilization of the leaving group [9, 10]. These four principal catalytic strategies used to promote RNA scission by phosphoester transfer are depicted in Figure 1.

**FIG. 1.**
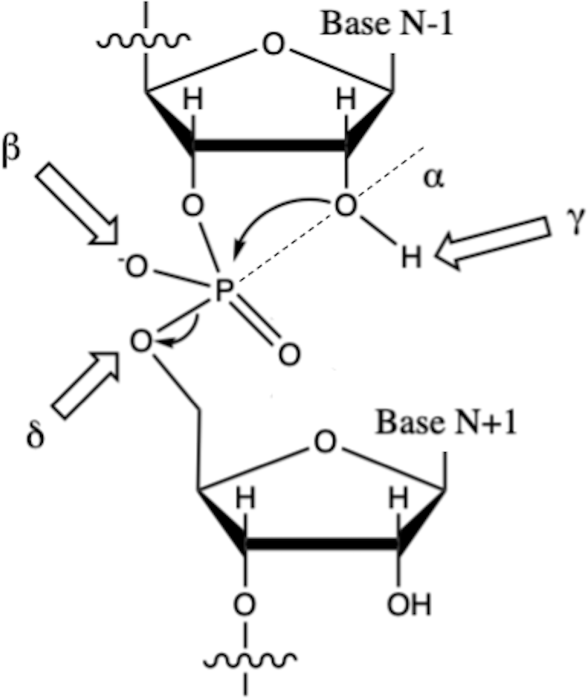
The mechanisms of the cleavage reaction between nucleotides N-1 and N+1. An in-line conformation, depicted by the *α* axis (the dotted line), facilitates the S_N_2 attack of the 2′-oxygen atom of the N-1 nucleotide on the phosphate center, liberating the N+1 nucleotide. The other strategies that can be utilized to accelerate this reaction are denoted as follows: *γ* (proton abstraction), *β* (neutralization of non-bridging phosphate oxygen atoms), and *δ* (stabilization of the developing negative charge on the 5′-oxygen atom).

In-line attack (*α* strategy) requires an axis running through the 2′-oxygen nucleophile, adjacent phosphorus atom and the 5′-oxygen leaving group. This establishes a crucial line along which the participating atoms must be appropriately aligned for an S_N_2-like process to occur [16, 17]. Enhancement of nucleophilic attack can be achieved by neutralizing the negative charge on non-bridging phosphate oxygen atoms (*β* strategy). Abstraction of the proton from the 2′-hydroxyl (*γ* strategy) further increases the nucleophilicity of the attacking oxygen, thereby enhancing catalysis. Similarly, neutralizing the developing charge on the 5′-oxygen leaving group (*δ* strategy) also promotes catalysis [9, 13, 18]. These mechanistic delineations of the cleavage reaction are crucial for understanding the activity of the molecule. Biochemical assays and atomic-resolution structural models support this simple framework for understanding the mechanism of catalysis [8, 9, 19–21].

Pistol ribozymes accelerate RNA catalysis via this internal phosphoester transfer mechanism with reaction rate constants as high as 10 min^−1^ under physiological conditions [19, 21]. The consensus sequence for pistol ribozymes was determined based on the alignment of 500 unique representatives found in DNA sequence databases comprised of complete bacterial genomes, microbial metagenomic DNA, and bacteriophage DNA [8].

Figure 2 depicts the three particular stems that comprise the secondary structure model of pistol ribozymes, referred to as P1, P2, and P3. Additionally, there is a pseudoknot predicted to form between the loop of P1 and the junction connecting P2 to P3.

**FIG. 2.**
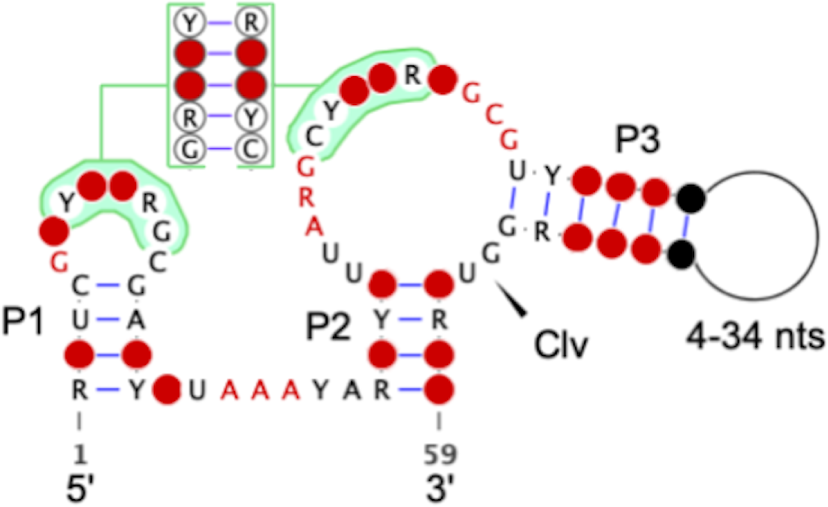
The consensus sequence and secondary structure model for pistol ribozymes. Red nucleotides are highly conserved, and the cleavage site is indicated by the arrowhead and occurs between guanosine and uridine nucleotides in our model RNAs. Several nucleotides (highlighted in green) interact to form a pseudoknot.

Crystal structures (PDB IDs 5K7C and 5KTJ) [20, 21] validate the aforementioned secondary structure [8, 19]. The presence of divalent cations may promote general acid-base catalysis and serve to coordinate with other critical residues in the pistol catalytic architecture [16]. However, the field lacks a comprehensive understanding of the influence of solvated divalent cations on catalysis. Recent work on twister ribozymes, and other self-cleaving RNAs, has shed light on the necessity of magnesium ions for maintaining structural integrity, as well as directly participating in catalysis [16, 22, 23]. In the current study on pistol ribozymes, we characterize the solvated divalent cations, which influence catalysis and distinguish them from the ions that influence conformational stability of the molecule.

Superficially, the presence of two magnesium ions in the vicinity of the S_N_2-like reaction center of pistol ribozymes (5K7C and 5KTJ) is reminiscent of the two-metal ion catalytic architecture prevalent among various enzymes [24]. However, catalysis driven by a single metal ion [25], as well as multiple metal ions [26], have also previously been characterized. These provide examples of metal-ion catalysis that are distinct from the previously observed two-metal mechanism. Hence, we closely follow the magnesium ions during the course of our simulations to understand their preferential participation in achieving near-catalytic states. Our principle focus in this study involved monitoring the magnesium ions that maintained close proximity to the catalytic site and/or moved from a starkly different position to a new coordination shell that can most likely influence catalysis.

## II. METHODS

In this study, several molecular dynamics simulations were performed using the 5K7C crystal structure of the pistol ribozyme (*env25* construct) as a starting model. The 5K7C structure model is based on the X-ray diffraction data yielding at 2.7 Å resolution and is comprised of an enzymatic chain of 47 residues and a substrate chain of 11 residues. The construct contains a deoxyribonucleotide at the cleavage site, ensuring that the observed crystal structure is not occluded by a hybrid of pre- and post-catalytic states. Additionally, three magnesium ions are present within this starting structure. Simulations were also performed with the analogous 5KTJ crystal structure for validation purposes. The 5KTJ structure contains several cobalt hexammine (III) complexes, which are expected to behave similar to other divalent metal ions in their solvated state due to the presence of coordinated amines [27]. We substituted each of the cobalt hexammine complexes with Mg^2+^ ions at the Co^2+^ site and associated water molecules to mimic the simulation conditions of the 5K7C model. In a previously reported study, a Mg^2+^ cation in close proximity to the reaction center was attributed solely to electrostatic stabilization rather than coordination with the non-bridging phosphate oxygen atoms [21].

For the purpose of simulations, the two chains of 5K7C were concatenated using a simple phosphodiester linker. Hydrogens were added to the original crystal structure using the `reduce` function of the PHENIX software [28]. We changed the hydrogen present at the 2′ position in the original structure to a hydroxyl group and ensured that the rotamer geometry was optimized using Coot software [29]. Upon addition of hydrogen atoms, we attained a refined model suitable for molecular dynamic simulations. This model for 5K7C was referred to as the “wild-type” (WT) model and will be referred to as such for the remainder of this article. This initial structure was immersed in a cubic box containing extended simple point charge model (SPC/E) water molecules and neutralized by addition of an appropriate number of sodium ions [30]. The ribozyme was centered in the box and placed at least 10 Å away from each edge. We also performed simulations using several other systemic parameters including SPC-E water model, TIP3P water model and CHARMM36 force field (Supplementary Figure 3), to exclude any possibility of building artifacts due to the choice of force field. However, we never observed any significant differences in the data leading to our postulates. A total of 0.921 *µ*s of simulation were performed on various WT and *in-silico* “mutants” in order to characterize the structural contribution of the three magnesium ions. In contrast to the aforementioned WT model, these mutants lack one or more magnesium ions or contain a substitution of sodium ions. The simulations in which certain magnesium ions were removed will be referred to as “dropouts” and will be denoted with a Δ for the remainder of this report (Table 1).

**TABLE I.** The various classical force field MD simulations performed. Various Mg^2+^ ions were present in each structure to probe their effect on overall conformational stability and their potential to influence catalytic geometry. Each structure was energy minimized, and subsequently relaxed in NVT, followed by NPT, conditions. At least five production-length trajectories of 10ns were concatenated to yield a minimum total of roughly 50ns of trajectory data for each mutant. All simulations were carried out with the CHARMM27 all-atom force field in Gromacs.

| Name | PDB | Description | Time |
| --- | --- | --- | --- |
| WT | 5K7C | Contains all three $\text{Mg}^{2+}$ ions present in the 5K7C structure. | 219 ns |
| WT | 5KTJ | Contains both original $\text{Mg}^{2+}$ in the 5KTJ structure, as well artificially inserted $\text{Mg}^{2+}$ ions that replace the original cobalt hexamines. | 50 ns |
| $\Delta\text{MG101}$ | 5K7C | Removal of MG101, located 13.9 Å from the phosphate center. | 40 ns |
| $\Delta\text{MG102}$ | 5K7C | Removal of MG102, located 5.0 Å from the phosphate center. | 205 ns |
| $\Delta\text{MG103}$ | 5K7C | Removal of MG103, located 7.8 Å from the phosphate center. | 203 ns |
| $\Delta\text{MG102}\Delta\text{MG103}$ | 5K7C | Only MG101 present. | 81 ns |
| MG101toNA | 5K7C | Sodium substitution at position of MG101 | 50 ns |
| MG102toNA | 5K7C | Sodium substitution at position of MG101 | 50 ns |
| MG103toNA | 5K7C | Sodium substitution at position of MG101 | 50 ns |
| MG101toNA, $\Delta\text{MG102}\Delta\text{MG103}$ | 5K7C | Sodium substitution at position of MG101 and removal of MG102 and MG103 | 50 ns |

In addition to 5K7C mutants, we also performed analogous dropout simulations using the 5KTJ structure, which was crystallized along with several cobalt hexammine coordination complexes. These complexes were replaced with hydrated magnesium ions at the position of Co^2+^ for the purposes of simulation.

All simulations were performed with the `mdrun` sub-routine of Gromacs with the CHARMM27 all-atom force field [31, 32]. Each system was initially energy minimized. Energy minimization was performed using the steepest descent minimization integrator, a Verlet cuttoff-scheme [33], the Particle Mesh Ewald (PME) method for electrostatic interactions [34], and periodic boundary conditions. After minimization, the system was relaxed under NVT conditions and NPT conditions at 300 K with isotropic Parrinello-Rahman pressure coupling [35]. This relaxation process used the leap-frog integrator, a `dt` of 2 fs, PME, and Verlet cutoff. Production MD runs were performed using the aforementioned settings, however for longer periods of time.

We also performed test runs with AMBER14 force field, CHARMM36 force field, and the TIP3P water model (data not shown). We did not find any significant differences in our mechanistic model, especially pertaining to the WT and magnesium dropout trajectories.

Trajectory analysis was performed using the standard `sci-py` stack [36] and the `mdanalysis` Python library [37, 38]. Data analysis was performed in an `iPython jupyter` notebook environment using the `numpy` and `pandas` packages [39–42]. Molecular visualizations were generated using the `nglview` package and `PyMol` software [43–46]. Plots and figures were generated with the `matplotlib` and `seaborn` package [47, 48].

RMSDs for each frame of the trajectory were calculated and binned to create RMSD distributons for various *in-silico* mutants according to the following equation [49]

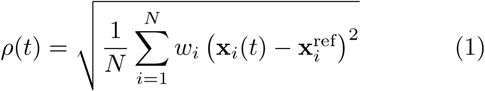

where *ρ*(*t*) is the time-dependent RMSD, *N* is the number of atoms, **x**_*i*_(*t*) is the vector of atomic coordinates for the *i*th residue at time *t*, and 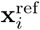 is the vector of atomic coordinates of the *i*th residue of the minimized and relaxed structure. Time-dependent RMSD fluctuations over the course of simulation can be considered a metric of overall stability and conformational rigidity.

Additionally, pair-distribution functions, known as radial distribution functions (RDFs) were calculated for each magnesium ion with respect to the catalytic phophate center. The RDF effectively counts the average number of *b* particles (a specific magnesium ion) in a shell at distance *r* around an *a* particle (the phosphate center) and represents it as a density (assuming particle *a* was chosen as the origin of the coordinate system). The RDF *g*_*ab*_(*r*) can be represented as

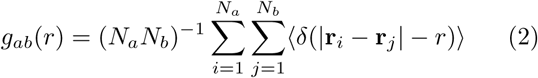

where *N*_*a*_ and *N*_*b*_ denote the number of particles of type *a* and type *b, i* and *j* are indexing variables used in the summation, **r**_*i*_ and **r**_*j*_ are vectors to the *i*th and *j*th particles respectively [50]. This above equation simplifies when only one of each particle type is present:

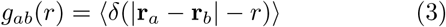

Upon inspection, it becomes clear that Equation 3 provides the time averaged frequency of finding particles *a* and *b* at a distance *r* from each other. Since our analysis, for the most part, involved analyzing groups of particles in which there was only one member molecule/atom, the RDF plots obtained were analogous to Euclidean distance distributions. Furthermore, over the course of the trajectories, the magnesium ions of interest near the catalytic site maintained a complexed state with the macromolecule. In our case, the magnesium ions remained at a significant distance from the phosphate center and both the bridging and non-bridging oxygen atoms, allowing us to rule out the possibility of a direct coordination and instead favor a solvated interaction.

## III. RESULTS & DISCUSSION

The structures of the *env25* pistol ribozyme was analyzed in their equilibrium states in several different contexts (WT and Δs of magnesium ions, see *Methods* section) in neutralizing aqueous environments. The reference PDB 5K7C used for the study was modified to contain the usual RNA nucleotide at the cleavage site instead of its DNA counterpart. The primary hypothesis of the study is that the presence of the native ribonucleotide allows the RNA molecule to achieve conformations that are closer to its usual catalytic states. The overall RMSD fluctuations were within the range of ± 1 Å for all the trajectories. Larger fluctuations were observed in trajectories containing monovalent sodium substitutions of the corresponding divalent ions.

Production MD runs on the 5K7C variants, including the magnesium dropouts (Δs), displayed interesting dynamic properties. The dropout of each of the magnesium atoms from the simulations yielded very small differences in the RMSD distributions. The width of the distributions remained relatively constant, but the mean values changed in some cases, as depicted in Figure 4.

**FIG. 3.**
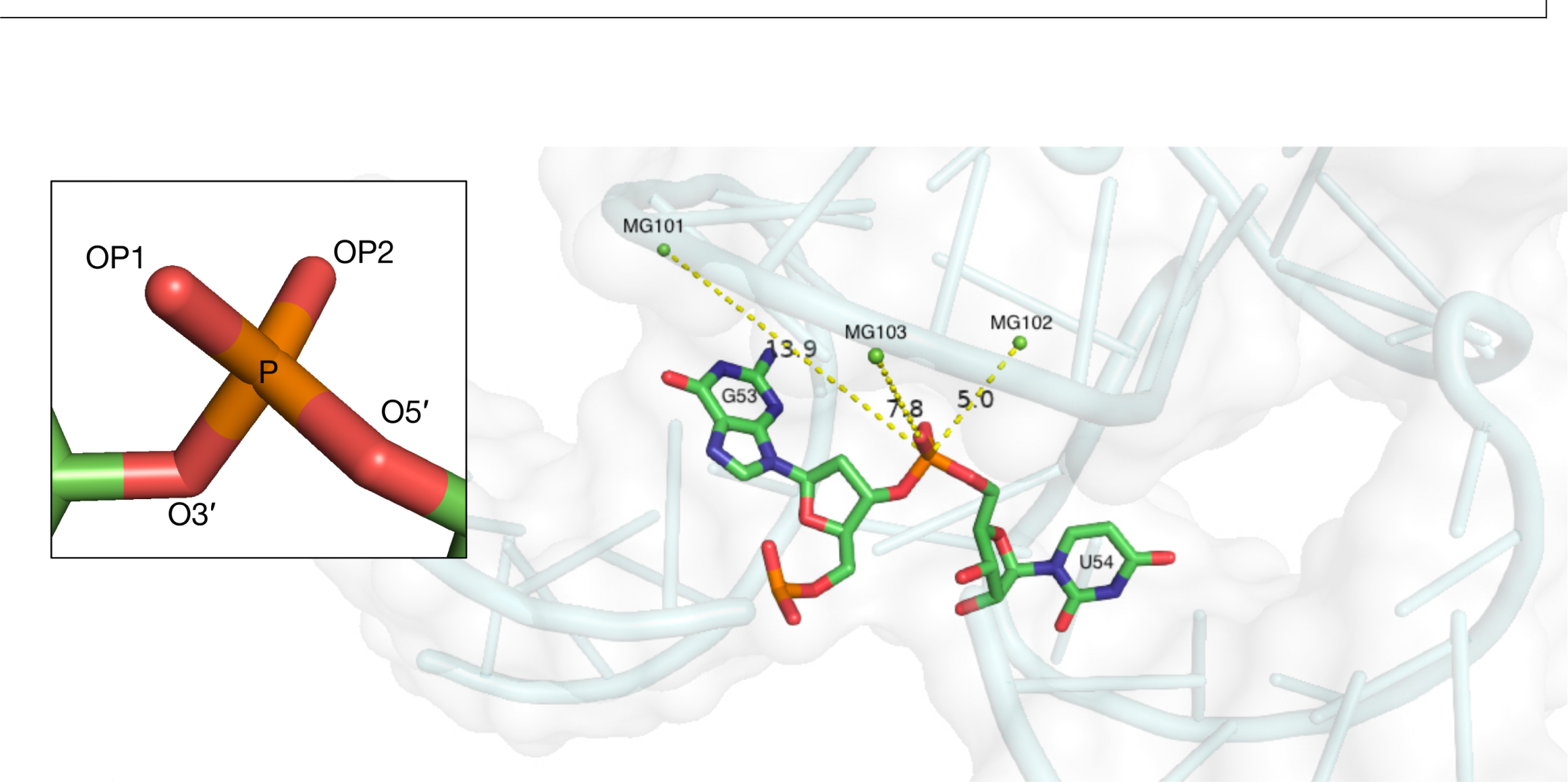
5K7C crystal structure with the the dG and U residues that form the cleavage site emphasized, including a view of the phosphate reaction center. The cleavage site residues are depicted as sticks, and the RNA is represented with gray space-filling spheres and cartoon backbone. The three magnesium ions present in the structure are shown as green spheres and labeled with distances to the phosphate center indicated in Å. MG102 and MG103 are close enough to the phosphate center to potentially play a role in the S_N_2 reaction [21]. The inset depicts the phosphate center with non-bridging oxygen atoms OP1 (*pro*-S) and OP2 (*pro*-R), and bridging oxygen atoms O5′ and O3′.

**FIG. 4.**
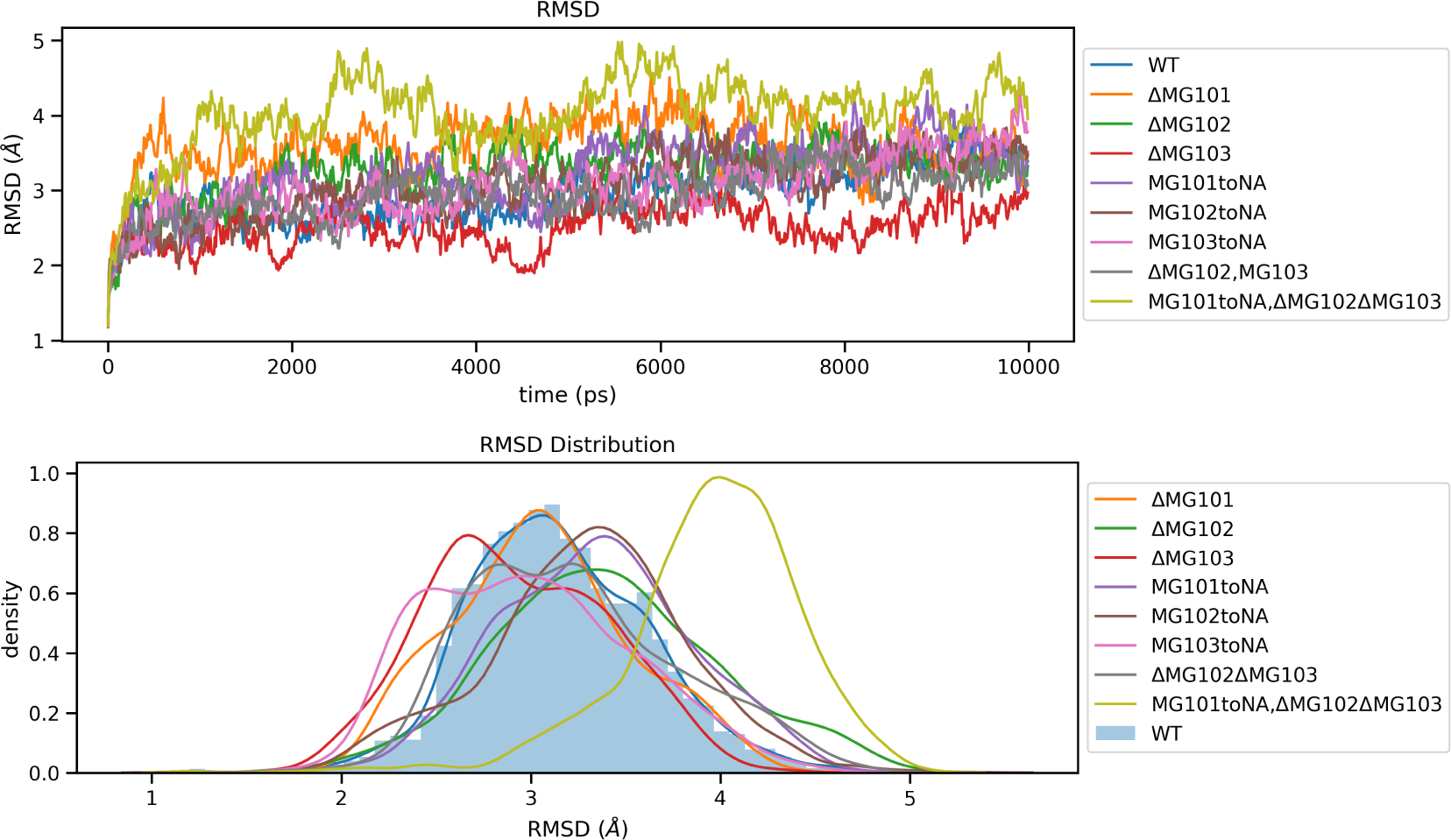
(Top) RMSDs from relaxed 5K7C structures during a 10 ns trajectory propagation. (Bottom) Distribution of RMSDs of 5K7C structures over 50 ns period.

The RMSD fluctuations in the dropout simulations were also larger than the WT simulations but smaller than their sodium-substitution counterparts. It is interesting to note that the sodium replacement of MG101 alongside the removal of MG102 and MG103 had the most effect on the overall RMSDs. The radius of gyration of the entire molecule remained stable throughout the course of the WT and magnesium-dropout simulations, hinting towards a consistent compactness of the RNA molecule (Figure 5).

**FIG. 5.**
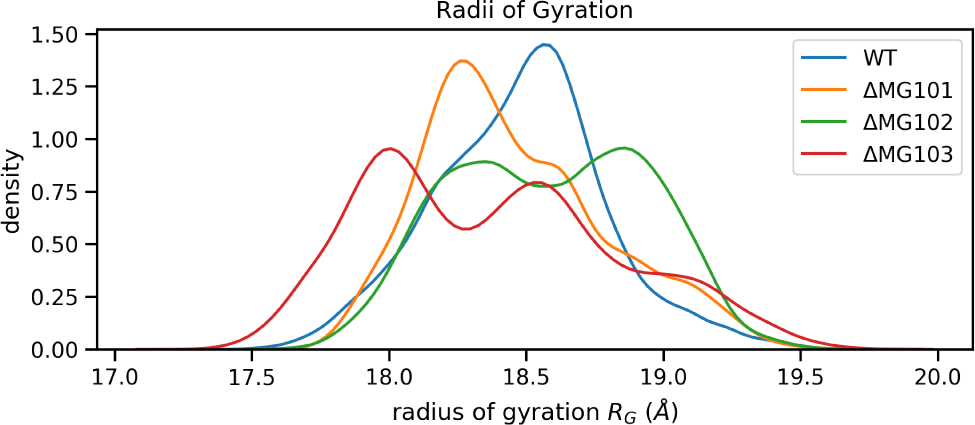
Radius of gyration during simulation of the 5K7C variants.

During the trajectories of every simulation variant, the MG101 atom populated a distal site at least 11 Å away from the phosphorous center, as seen in every radial distribution function plot in Figure 6. The magnesium-dropout, as well as sodium-substitution, at position 101 did not affect the overall structure of the molecule. However, a substitution with sodium in that same position without any other magnesium ions in the system led to a higher mean RMSD of the molecule by roughly 1 Å. When the simulations with the 5K7C structure were compared to the simulations with the 5KTJ structure, the results hold a similar pattern of change. The overall increase in the RMSD of the molecule, coupled with the distant nature of MG101 from the catalytic site, strongly suggest that this ion position primarily plays a structural role.

**FIG. 6.**
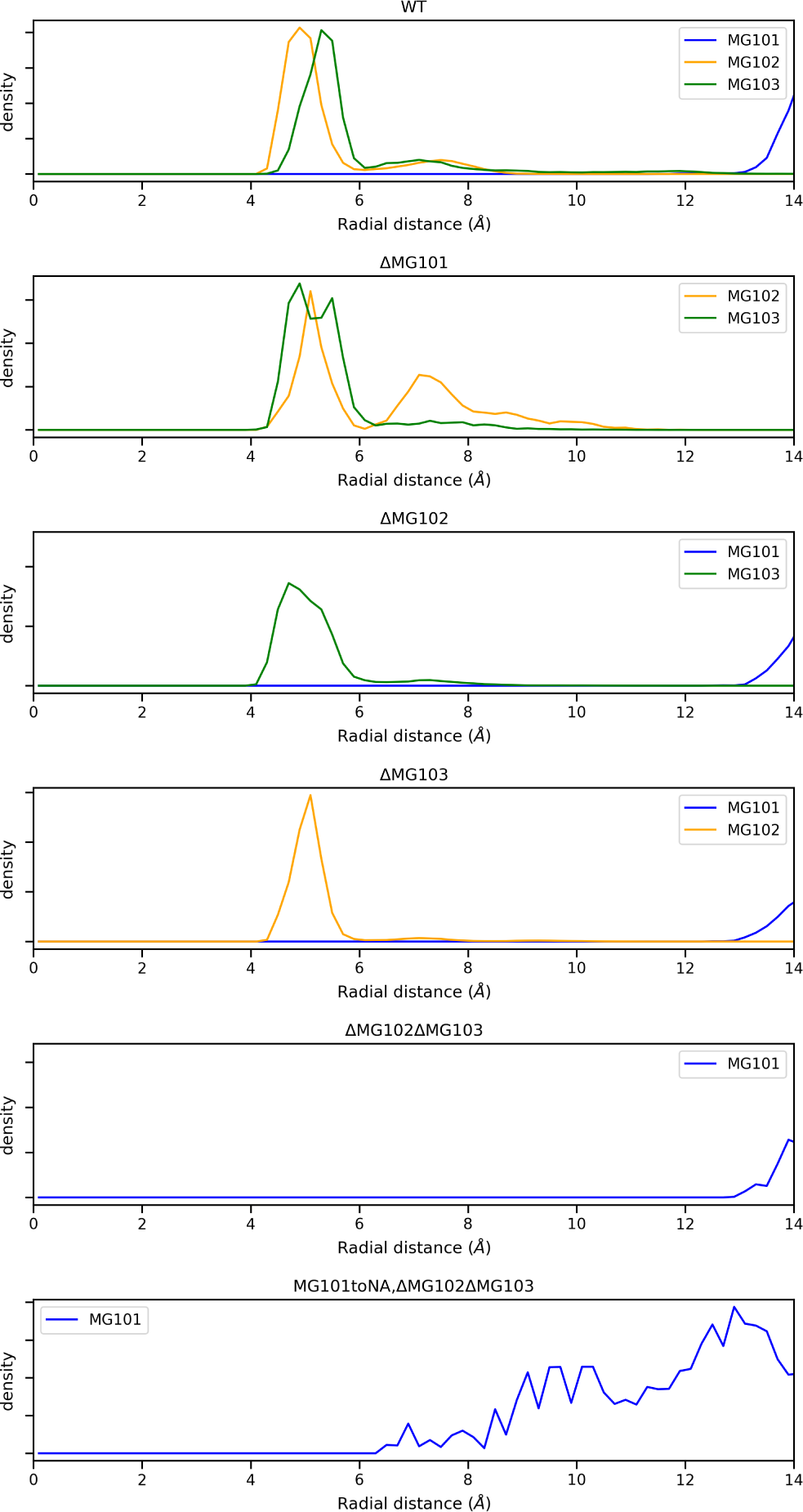
Radial distributions between the phosphate center and MG101, MG102, and MG103 for various 5K7C *in-silico* mutants.

It was also seen that ΔMG102 and ΔMG103 exhibited similar characteristics to the MG102toNA and MG103toNA simulations, respectively. The RMSD values for both of these conditions were within 1 Å of the mean RMSD of the wild-type simulation.

In 5K7C trajectories MG102, which existed in two populations (at 7 Å and at 4.5 Å) in the wild-type scenario, funneled completely into a single population (at 4.5 Å) in the case of the MG103 dropout simulation. An analogous funneling effect occurs in the 5KTJ trajectories. This result indicates a substantial contribution from MG102 towards stabilizing the catalytic site in the absence of MG103. The magnesium ions close to the catalytic center preferentially populate a proximal site capable of coordinating solvated hydrogen bonds rather than a distal site as depicted in the crystal structures [20, 21].

Systematic characterization of each of the 14 solvated cations from the 5KTJ structure, provided a comprehensive picture of their localization and their plausible roles in catalysis. MG101, MG104, MG105, MG106, MG107, MG108, MG109, MG110, MG111, MG112, MG113, and MG114 were all localized far from the catalytic core of the ribozyme during the entire course of the simulations. MG102 and MG103 which were crucial in the case of 5K7C also localized within solvated hydrogen bonding distance from the catalytic core in the case of the 5KTJ simulations (*Supplementary* Figure 1).

We wanted to take a closer look at these two magnesium ions (MG102 and MG103) and also wanted to characterize their interactions with the oxygen atoms, which are likely to be involved in catalysis. The top two plots of Figure 7 depict the distance from the magnesium ions to each of the oxygen atom at the cleavage site that are involved in attaining the catalytic geometry of the molecule. The distance from every oxygen atom to MG102 remains predominantly constant in the MG103 dropout simulations, similar to the WT scenario.

**FIG. 7.**
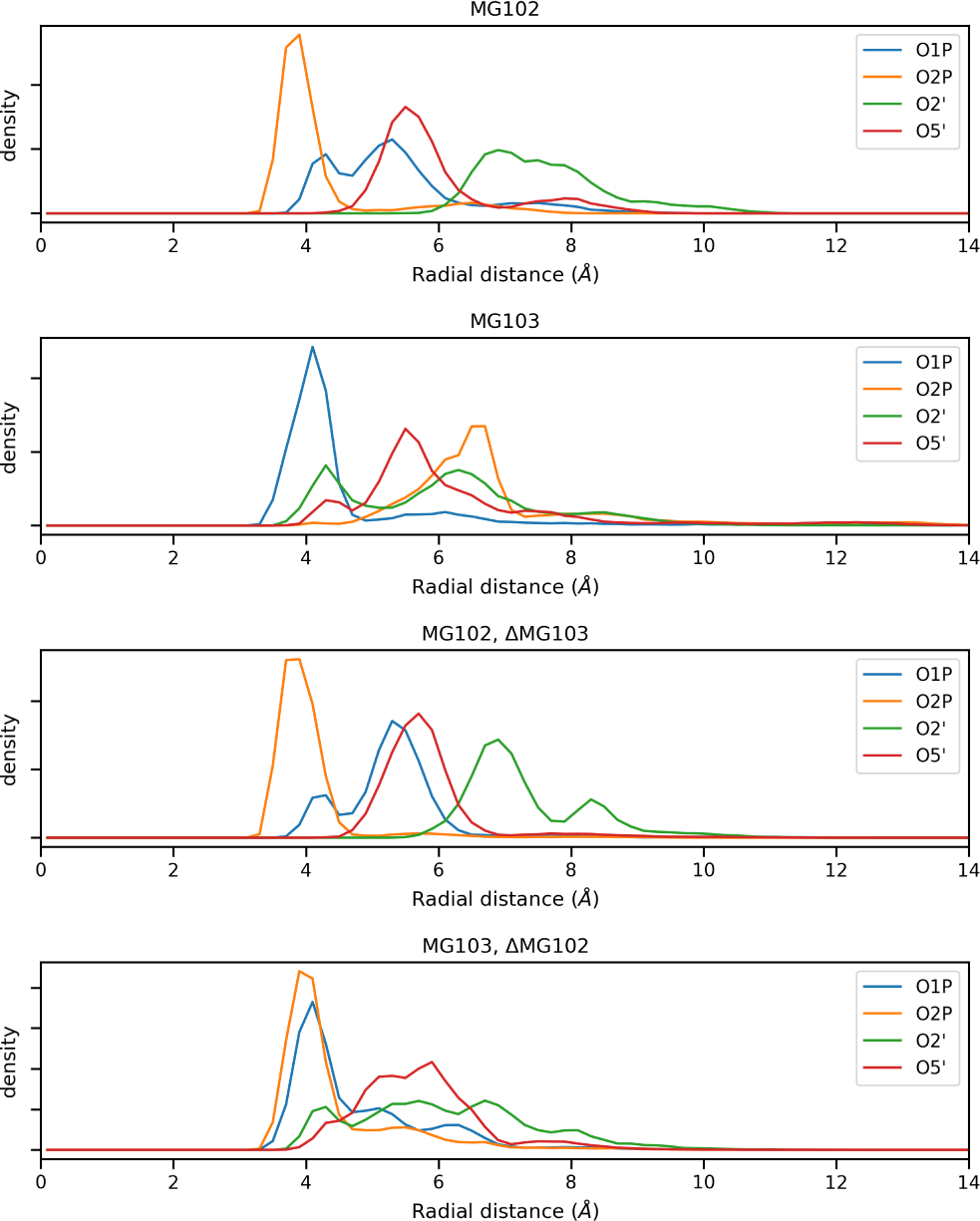
Radial distributions of magnesium ions and relevant sites on the phosphate center in the 5K7C WT and mutant simulations.

When comparing the bottom two panels of Figure 7, we see a stark shift in the occupancy of the 4 Å site. In the ΔMG102 trajectories, magnesium at the 103 site diffuses closer to the catalytic center, stabilizing the coordination of the O2P non-bridging oxygen (*pro*-R) at the cost of the 5′-oxygen attached to the phosphorous center. This indicates a preferential stabilization of the non-bridging oxygen atoms by two separate magnesium ions in WT environment; O2P with MG102 and O1P with MG103. However, in case of the MG102 dropout, MG103 takes on the additional role of coordinating with O2P, thereby providing a fail-safe mode for catalysis (Schematic in *Supplementary* Figure 2).

When we probed the distributions of the approach angle of the transition state between the 2′-oxygen and the phosphorus center at the cleavage site, we noticed a range of conformations being sampled by the molecule, peaking around 135 degrees. It is conceivable that the closer the molecule samples an angle of 180 degrees, the closer it is to a transition state and hence favourably undergoes the cleavage reaction. We can see from Figure 8 that even though MG101 does not play any direct role in coordinating with the active site, its deletion pushes the equilibrium to a secondary unfavourable population.

**FIG. 8.**
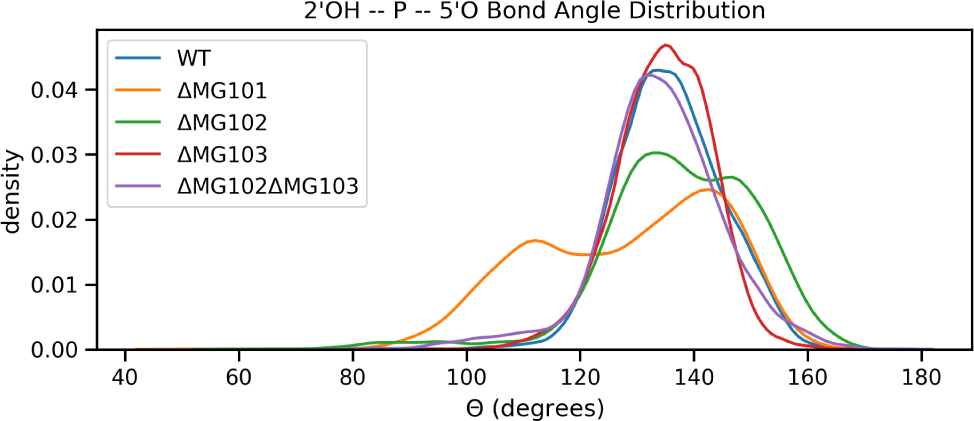
Bond angle distribution in the various 5K7C structure. Bond angles closer to 180 degrees are more favorable for the S_N_2 in-line attack of the 2′-OH of the N-1 nucleotide on the phosphorus of the N+1 nucleotide.

It is also intriguing that even in the absence of magnesium ions near the catalytic site (as in the ΔMG102 and ΔMG103 environments), MG101 still maintains its position, and hence could be inferred to play only a structural role in the folding of this molecule (Figure 6). It is far too distant to be directly involved at the active site of the RNA. Nevertheless, maintaining the proper fold of the RNA is also important for the overall ability to attain the catalytic conformations (Figure 7).

## IV. SIGNIFICANCE

Our *in-silico* results are consistent with the hypothesis that solvated magnesium ions play a crucial role in catalysis by pistol ribozymes. This conclusion is in close accordance with the previous mutational studies. These ribozymes, in part, employ *β*-catalysis for cleavage activity, which diminishes drastically in the absence of divalent magnesium ions [51]. Previous work on atom-specific mutagenesis of Pistol ribozyme indicated an impactful role of a specific Adenosine (32nd in *env25*) along with the necessity of a hydrated Magnesium ion. There was a strong suggestion from the same article that a water molecule linked to the Magnesium ion probably acts as a proton donor and the stabilization of the non-bridging oxygen is crucial for accelerating the process [51]. While this manuscript was under preparation a detailed molecular dynamics study was reported by Kostenbader and co-workers, which provided critical insights into the similarities between pistol and hammerhead ribozymes. This enables the community to look beyond the specific intricacies of individual systems and address the broader evolutionary picture consisting of all ribozymes. Koestenbader *et. al.* dives deep into examining the nucleobase organization at the active site and its influence in catalysis [52]. The focus of this recent publication is quite complementary to our work as we focused on understanding the distribution of divalent cations and their effects on pre-catalytic states. In the article the authors also comment on how the binding of these Magnesium ions are subtly different in the two ribozyme systems they have compared. It is curious that the rescue effect of Magnesium movements highlighted in the article (for hammerhead ribozyme)[53], where the Magnesium moves from a non-bridging position to a bridging position, is observed to be completely recapitulated in our pistol ribozyme simulations. Our results are in close accordance with the hypothesis of Pro-R oxygen stabilization while also proving the importance of the cation interaction site. Additionally, these simulations stipulate that the 102 divalent position is crucial for catalysis, such that removal of the magnesium ion from that site induces another magnesium ion to assume its position. These findings provide computational plausibility for a solvated magnesium-dependent mechanism of self-cleavage employed by pistol ribozymes. It consolidates the existing understanding of the influence of divalent cations on the various catalytic strategies, while suggesting the existence of a fail-safe secondary metal ion. These types of arrangements in catalytic RNA hint towards more sophisticated functions of divalent cations, along with their malleability in non-uniform structural and dynamic catalytic roles. A thorough study with the characterization of transition-state analogue along with a time-dependant cation analysis will further solidify our claim. Our work resonates with the current paradigm of catalytic mechanism in pistol ribozymes while describing the novel facet of the ionic contribution. This type of catalytic architecture could very well be true for many other cation dependent reactions. Even though we comparatively examine roles of magnesium ions associated with the structures of pistol ribozymes, further work is necessary to experimentally validate the concept of fail-safe ions and quantify their individual effects on mechanisms of self-cleaving ribozymes.

## ACKNOWLEDGMENTS

We would like to thank the Yale Center for Research Computing for computational time and access to the Grace cluster. Additionally, we thank the staff of the Richards Center at Yale University for technical support. We are also thankful to Professor Ronald R. Breaker for advice and critical readings of the manuscript. We are thankful to Zachary Levine for his help in assisting us through simulation-related technical matters.

We thank the Howard Hughes Medical Institute and the National Institutes of Health [GM022778 to T.A.S.] for funding.

We are deeply thankful to Professor Thomas A. Steitz for his guidance, insight, and mentorship during our time in his lab. Despite his passing, his impact on his students and the field as a whole is indelible.

R.N.R. and N.N.J. performed experiments, analyzed data, and wrote manuscript as joint-first authors.

## V. SUPPLEMENTARY INFORMATION

**FIG. 1.**
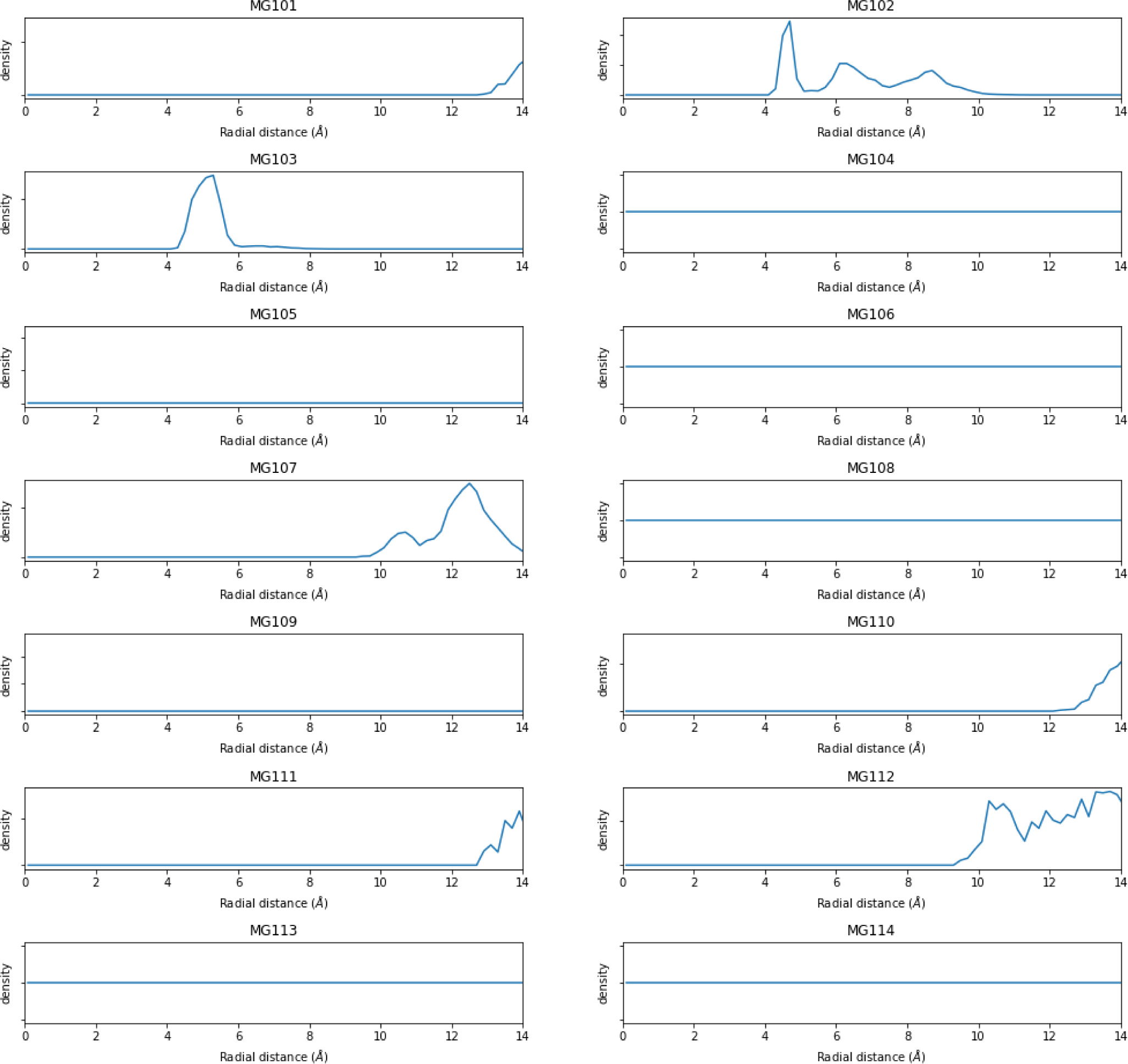
Radial distributions between magnesium ions and the phosphate center in the 5KTJ WT simulation.

**FIG. 2.**
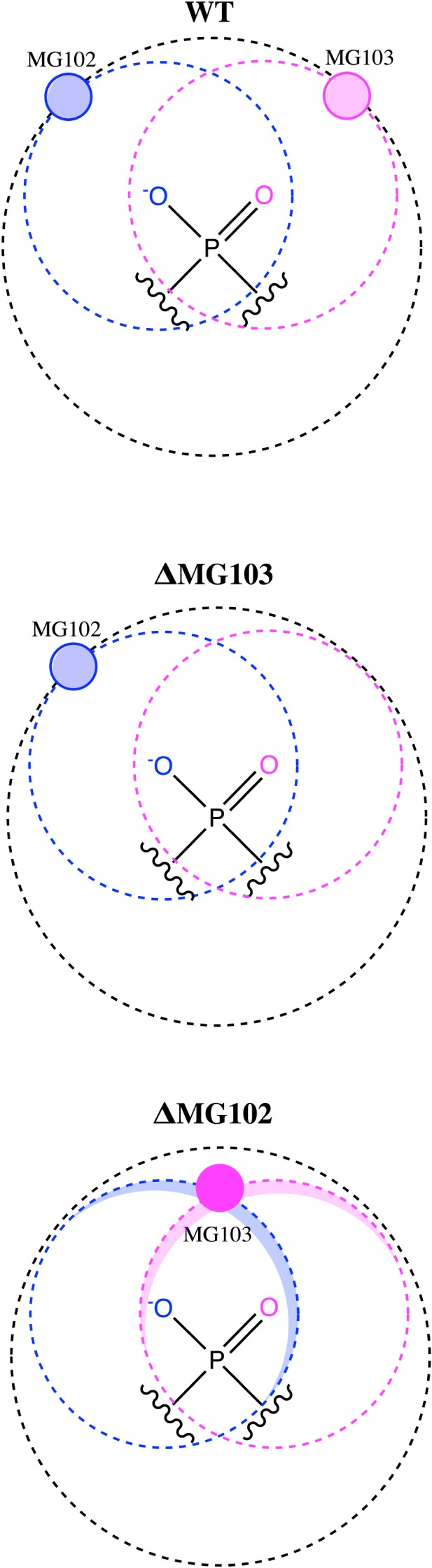
Schematic representation of the movements of the Mg^2+^ ions. The top panel highlights the 4.5 Å shell from the phosphorus center at which MG102 and MG103 preferentially resides (black circle). The blue and the pink shells represent their individual distances from the two non-bridging oxygen atoms. The middle panel shows that MG102 maintains its distances from the phosphorus and its oxygen partner in absence of MG103. In the bottom panel, MG103 goes out of the plane (dark red) to move closer to the non-bridging *pro*-R oxygen (blue) while maintaining its distance from the phosphorus center.

**FIG. 3.**
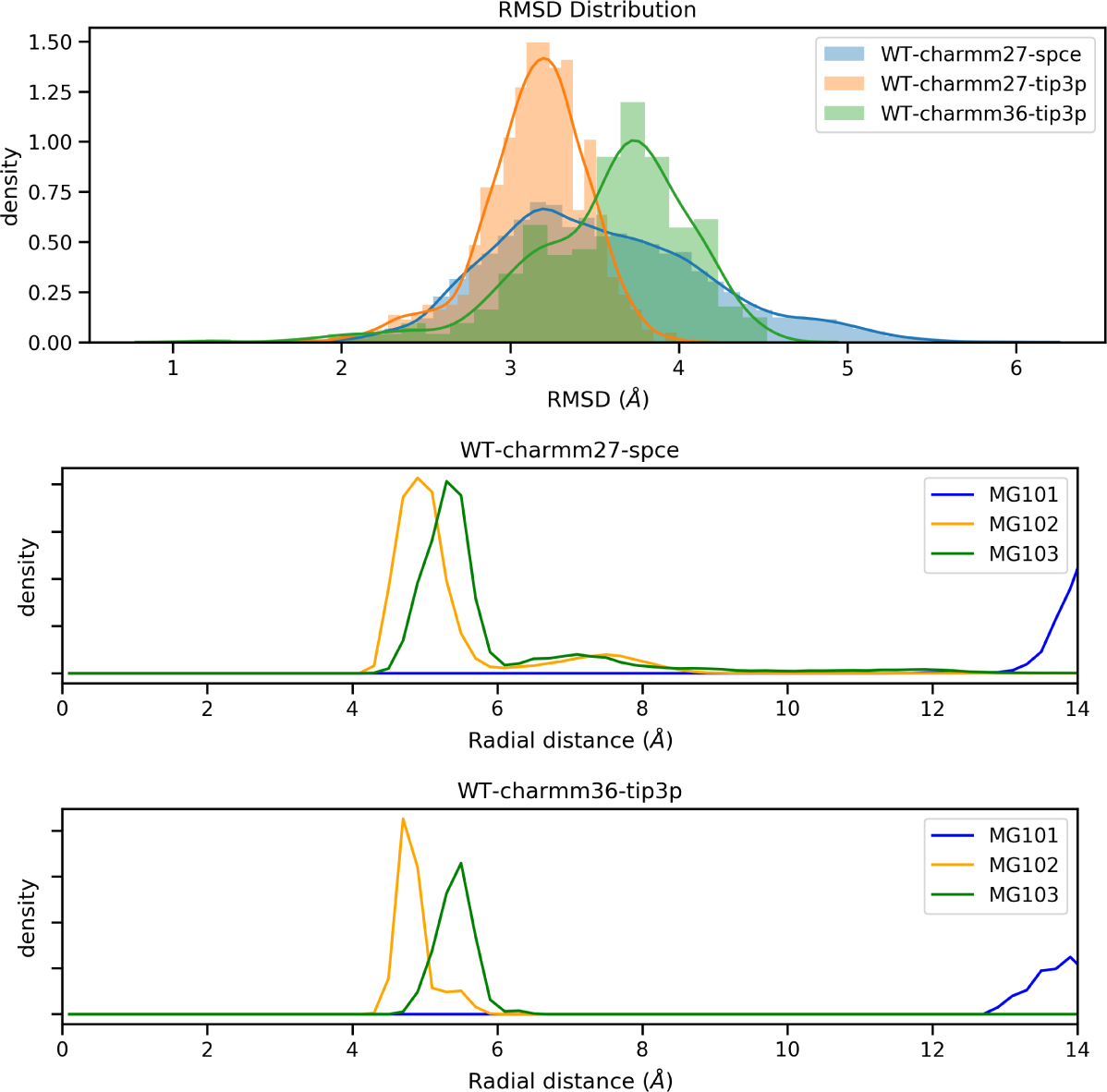
RMSD and RDF distributions across different systemic parameters. The top panel highlights the shows the normalized profiles of RMSD distributions in different water models and force fields. In the subsequent panels, RDF distributions of magnesium ions from the catalytic phosphate center are shown.

## REFERENCES

[1] Prody, G. A., Bakos, J. T., Buzayan, J. M., Schneider, I. R., and Bruening, G. (1986) Autolytic processing of dimeric plant virus satellite RNA. Science 231, 1577–1580.

[2] Hutchins, C. J., Rathjen, P. D., Forster, A. C., and Symons, R. H. (1986) Self-cleavage of plus and minus RNA transcripts of avocado sunblotch viroid. Nucleic Acids Res. 14, 3627–3640.

[3] Buzayan, J. M., Gerlach, W. L., and Bruening, G. (1986) Non-enzymatic cleavage and ligation of RNAs complementary to a plant virus satellite RNA. Nature 323, 349.

[4] Kuo, M., Sharmeen, L., Dinter-Gottlieb, G., and Taylor, J. (1988) Characterization of self-cleaving RNA sequences on the genome and antigenome of human hepatitis delta virus. J. Virol. 62, 4439–4444.

[5] Saville, B. J., and Collins, R. A. (1990) A site-specific self-cleavage reaction performed by a novel RNA in Neurospora mitochondria. Cell 61, 685–696.

[6] Winkler, W. C., Nahvi, A., Roth, A., Collins, J. A., and Breaker, R. R. (2004) Control of gene expression by a natural metabolite-responsive ribozyme. Nature 428, 281–286.

[7] Roth, A., Weinberg, Z., Chen, A. G., Kim, P. B., Ames, T. D., and Breaker, R. R. (2014) A widespread self-cleaving ribozyme class is revealed by bioinformatics. Nat. Chem. Biol. 10, 56–60.

[8] Weinberg, Z., Kim, P. B., Chen, T. H., Li, S., Harris, K. A., Lünse, C. E., and Breaker, R. R. (2015) New classes of self-cleaving ribozymes revealed by comparative genomics analysis. Nat. Chem. Biol. 11, 606–610.

[9] Breaker, R. R. (2017) Mechanistic debris generated by twister ribozymes. ACS Chem. Biol. 12, 886–891.

[10] Emilsson, G. M., Nakamura, S., Roth, A., and Breaker, R. R. (2003) Ribozyme speed limits. RNA 9, 907–918.

[11] Ferré-D’Amaré, A. R., and Scott, W. G. (2010) Small self-cleaving ribozymes. Cold Spring Harb. Perspect. Biol. 2, a003574.

[12] Lilley, D. M. (2011) Mechanisms of RNA catalysis. Philos. Trans. R. Soc. Lond. B: Biol. Sci. 366, 2910–2917.

[13] Doherty, E. A., and Doudna, J. A. (2000) Ribozyme structures and mechanisms. Ann. Rev. Biochem. 69, 597–615.

[14] Breaker, R. R., Emilsson, G. M., Lazarev, D., Nakamura, S., Puskarz, I. J., Roth, A., and Sudarsan, N. (2003) A common speed limit for RNA-cleaving ribozymes and deoxyribozymes. RNA 9, 949–957.

[15] Wilson, T. J., and Lilley, D. M. (2009) The evolution of ribozyme chemistry. Science 323, 1436–1438.

[16] Liu, Y., Wilson, T. J., McPhee, S. A., and Lilley, D. M. (2014) Crystal structure and mechanistic investigation of the twister ribozyme. Nat. Chem. Biol. 10, 739.

[17] Soukup, G. A., and Breaker, R. R. (1999) Relationship between internucleotide linkage geometry and the stability of RNA. RNA 5, 1308–1325.

[18] Lee, K.-Y., and Lee, B.-J. (2017) Structural and Biochemical Properties of Novel Self-Cleaving Ribozymes. Molecules 22.

[19] Harris, K. A., Lünse, C. E., Li, S., Brewer, K. I., and Breaker, R. R. (2015) Biochemical analysis of pistol self-cleaving ribozymes. RNA 21, 1852–1858.

[20] Nguyen, L. A., Wang, J., and Steitz, T. A. (2017) Crystal structure of Pistol, a class of self-cleaving ribozyme. Proc. Natl. Acad. Sci. U.S.A. 114, 1021–1026.

[21] Ren, A., Vušurović, N., Gebetsberger, J., Gao, P., Juen, M., Kreutz, C., Micura, R., and Patel, D. J. (2016) Pistol ribozyme adopts a pseudoknot fold facilitating site-specific in-line cleavage. Nat. Chem. Biol. 12, 702.

[22] Ucisik, M. N., Bevilacqua, P. C., and Hammes-Schiffer, S. (2016) Molecular dynamics study of twister ribozyme: role of Mg2+ ions and the hydrogen-bonding network in the active site. Biochemistry 55, 3834–3846.

[23] Murray, J. B., Seyhan, A. A., Walter, N. G., Burke, J. M., and Scott, W. G. (1998) The hammerhead, hairpin and VS ribozymes are catalytically proficient in monovalent cations alone. Chem. Biol. 5, 587–595.

[24] Steitz, T. A., and Steitz, J. A. (1993) A general two-metal-ion mechanism for catalytic RNA. Proc. Natl. Acad. Sci. U.S.A. 90, 6498–6502.

[25] Hu, X., Machius, M., and Yang, W. (2003) Monovalent cation dependence and preference of GHKL ATPases and kinases 1. FEBS Lett. 544, 268–273.

[26] Gao, Y., and Yang, W. (2016) Capture of a third Mg2+ is essential for catalyzing DNA synthesis. Science 352, 1334–1337.

[27] Cowan, J. (1993) Metallobiochemistry of RNA. Co (NH3) 63+ as a probe for Mg2+ (aq) binding sites. J. Inorg. Biochem. 49, 171–175.

[28] Adams, P. D., Afonine, P. V., Bunkόczi, G., Chen, V. B., Davis, I. W., Echols, N., Headd, J. J., Hung, L.-W., Kapral, G. J., Grosse-Kunstleve, W. R. (2010) PHENIX: a comprehensive Python-based system for macromolecular structure solution. Acta Crystallogr. Sect. D: Biol. Cryst. 66, 213–221.

[29] Emsley, P., Lohkamp, B., Scott, W. G., and Cowtan, K. (2010) Features and development of Coot. Acta Crystallogr. Sect. D: Biol. Cryst. 66, 486–501.

[30] Berendsen, H., Grigera, J., and Straatsma, T. (1987) The missing term in effective pair potentials. J. Phys. Chem. 91, 6269–6271.

[31] Abraham, M. J., Murtola, T., Schulz, R., Páll, S., Smith, J. C., Hess, B., and Lindahl, E. (2015) GROMACS: High performance molecular simulations through multi-level parallelism from laptops to supercomputers. SoftwareX 1, 19–25.

[32] Brooks, B. R., Brooks III, C. L., Mackerell Jr, A. D., Nilsson, L., Petrella, R. J., Roux, B., Won, Y., Archontis, G., Bartels, C., Boresch,, Stefan, (2009) CHARMM: the biomolecular simulation program. J. Comp. Chem. 30, 1545–1614.

[33] Páll, S., and Hess, B. (2013) A flexible algorithm for calculating pair interactions on SIMD architectures. Comp. Phys. Comm. 184, 2641–2650.

[34] Darden, T., York, D., and Pedersen, L. (1993) Particle mesh Ewald: An N log (N) method for Ewald sums in large systems. J. Chem. Phys. 98, 10089–10092.

[35] Fernández-Pendás, M., Escribano, B., Radivojević, T., and Akhmatskaya, E. (2014) Constant pressure hybrid Monte Carlo simulations in GROMACS. J. Mol. Model. 20, 2487.

[36] Jones, E., Oliphant, T., and Peterson, P. (2014) {SciPy} : Open source 5 scientific tools for Python. http://www.scipy.org. Online access.

[37] Michaud-Agrawal, N., Denning, E. J., Woolf, T. B., and Beckstein, O. (2011) MDAnalysis: a toolkit for the analysis of molecular dynamics simulations. J. Comp. Chem. 32, 2319–2327.

[38] Gowers, R. J., Linke, M., Barnoud, J., Reddy, T. J., Melo, M. N., Seyler, S. L., Dotson, D. L., Domanski, J., Buchoux, S., Kenney, M. I. Proceedings of the 15th Python in Science Conference; 2016; Vol. 98.

[39] Pérez, F., and Granger, B. E. (2007) IPython: a system for interactive scientific computing. Comput. Sci. Eng. 9, 21–29.

[40] Oliphant, T. E. A guide to NumPy; Trelgol Publishing USA, 2006; Vol. 1.

[41] Van Der Walt, S., Colbert, S. C., and Varoquaux, G. (2011) The NumPy array: a structure for efficient numerical computation. Comput. Sci. Eng. 13, 22.

[42] Mc Kinney,, Wes, Proc. 9th Python in Science Conf.; 2010; Vol. 445; pp 51–56.

[43] Rose, A. S., Bradley, A. R., Valasatava, Y., Duarte, J. M., Prlić, A., and Rose, P. W. NGL viewer: web-based molecular graphics for large complexes. 2018; pp 3755–3758.

[44] Nguyen, H., Case, D. A., and Rose, A. S. (2017) NGLview–interactive molecular graphics for Jupyter notebooks. Bioinformatics 34, 1241–1242.

[45] Tiemann, J. K., Guixa-Gonzalez, R., Hildebrand, P. W., and Rose, A. S. (2017) MDsrv: viewing and sharing molecular dynamics simulations on the web. Nat. Methods 14, 1123.

[46] Schrödinger, LLC, (2015) The PyMOL Molecular Graphics System, Version 1.8.

[47] Hunter, J. D. (2007) Matplotlib: A 2D graphics environment. Comput. Sci. Eng. 9, 90.

[48] Waskom, M., Botvinnik, O., OKane, D., Hobson, P., Lukauskas, S., and Gemperline, D. mwaskom/seaborn: v0. 8.1 2017. https://doi.org/10.5281/zenodo.883859.

[49] Theobald, D. L. (2005) Rapid calculation of RMSDs using a quaternion-based characteristic polynomial. Acta Crystallogr., Sect. A: Found. of Crystallogr. 61, 478–480.

[50] Van Leeuwen, J., Groeneveld, J., and De Boer, J. (1959) New method for the calculation of the pair correlation function. I. Physica 25, 792–808.

[51] Neuner, S., Falschlunger, C., Fuchs, E., Himmelstoss, M., Ren, A., Patel, D. J., and Micura, R. (2017) Atom-Specific Mutagenesis Reveals Structural and Catalytic Roles for an Active-Site Adenosine and Hydrated Mg2+ in Pistol Ribozymes. Angew. Chemie Intern. Ed. 56, 15954–15958.

[52] Kostenbader K, D. Y. (2019) Molecular simulations of the pistol ribozyme: unifying the interpretation of experimental data and establishing functional links with the hammerhead ribozyme. RNA

[53] Wang S, P. A. B. L. H. D., Karbstein K (1999) Identification of the Hammerhead Ribozyme Metal Ion Binding Site Responsible for Rescue of the Deleterious Effect of a Cleavage Site Phosphorothioate. Biochemistry 38, 14363–14378.

